# Age-related changes in spatial and temporal features of resting state fMRI

**DOI:** 10.1101/109181

**Authors:** Shruti G. Vij, Jason S. Nomi, Dina R. Dajani, Lucina Q. Uddin

## Abstract

Development and aging are associated with functional changes in the brain across the lifespan. These changes can be detected in spatial and temporal features of resting state functional MRI (rs-fMRI) data. Independent vector analysis (IVA) is a whole-brain multivariate approach that can be used to comprehensively assess these changes in spatial and temporal features. We present a multi-dimensional approach to assessing age-related changes in spatial and temporal features of statistically independent components identified by IVA in a cross-sectional lifespan sample (ages 6-85 years). We show that while large-scale brain network configurations remain consistent throughout the lifespan, changes continue to occur in both local organization and in the spectral composition of these functional networks. We show that the spatial extent of functional networks decreases with age, but with no significant change in the peak functional loci of these networks. Additionally, we show differential age-related patterns across the frequency spectrum; lower frequency correlations decrease across the lifespan whereas higher-frequency correlations increase. These changes indicate an increasing stability of networks with age. In addition to replicating results from previous studies, the current results uncover new aspects of functional brain network changes across the lifespan that are frequency band-dependent.

## Introduction

The human brain is composed of sub-sytems or networks that interact with each other forming a connectome (Sporns, Tononi et al. 2005, Kelly, Di Martino et al. 2009, Biswal, Mennes et al. 2010). These networks can be detected even in the absence of an external task, and present coherent communication patterns that can be characterized as resting-state networks (RSNs) or intrinsic connectivity networks (ICNs) (Biswal, Yetkin et al. 1995, Fox and Raichle 2007)(Seeley et al, 2007 J Neuro). Resting state networks such as the default mode network (DMN), the sensorimotor network (SMN), and others largely resemble networks activated during task performance (Smith, Fox et al. 2009). Recent advances in neuroimaging techniques allow for estimation or extraction of these ICNs using different methods such as independent component analysis (ICA) and its modifications and graph theoretical approaches (Calhoun and Adalı 2012, Betzel, Byrge et al. 2014, Hjelm, Calhoun et al. 2014) applied to the whole brain. While many studies have focused on changes in connectivity within and between ICNs (Geerligs, Renken et al. 2015, Huang, Hsieh et al. 2015), contemporary research exploring individual features such as the shape, size, and loci of ICNs and inter-subject variability of ICNs has been gaining importance (Grady and Garrett 2014, Laumann, Gordon et al. 2015, Yao, Palaniyappan et al. 2015).

Individual differences are widely known to influence individual features, in addition to interactions between networks (Jolles, van Buchem et al. 2011, Allen, Erhardt et al. 2012, Gordon, Laumann et al. 2016). A number of recent studies have evaluated age-related variability in some of these features, with within and between network connectivity being the primary focus (de Bie, Boersma M Fau - Adriaanse et al. 2012, Betzel, Byrge et al. 2014, Cao, Wang et al. 2014, Muetzel, Blanken et al. 2016, Sole-Padulles, Castro-Fornieles et al. 2016). These studieshave evaluated early development, adolescence to adulthood, as well as lifespan trajectories using both graph theoretical approaches and ICA based algorithms. The consensus from these studies is that ICNs are characterized by significant lifespan alterations in within-and between-network connectivity.

Other studies have focused on identifying neural substrates of behavior and cognitive maturation (Durston, Davidson et al. 2006, Fair, Cohen et al. 2009, Dickstein, Gorrostieta et al. 2010, Wang, Su et al. 2012, Bo, Lee et al. 2014). Cognitive maturation is the process by which problem solving, decision making, and other high-level processes become more refined with age(Khundrakpam, Lewis et al. 2016). Cognitive development and maturation are associated with network changes that manifest as modifications of ICN patterns as well as interactions between them. Earlier work using rs-fMRI demonstrated that while overall network structure is generally retained over lifespan, functional differences such as less spatial activation as well as connectivity changes are observed in ICNs involved in higher order cognitive abilities such as the DMN (Andrews-Hanna, Snyder et al. 2007, Fair, Cohen et al. 2009, Meunier, Achard et al. 2009, Cao, Wang et al. 2014, Huang, Hsieh et al. 2015). These studies verify that regional changes in ICNs are also widespread with age, and suggest that topological changes in ICNs can provide insights into neural substrates of individual variability (Jolles, van Buchem et al. 2011). Knowledge of this variability has important implications for understanding the development and maturation of higher-order cognitive abilities, and might be expressed as individual differences in cognitive strategies and/or behavioral differences.

Current advances in methods for the analysis of resting state fMRI data in conjunction with rising interest in identifying individual characteristics have led to examination of age-related changes in ICNs (Laumann, Gordon et al. 2015, Seghier and Price 2016). While thesestudies provide us with significant insight into the existence of variability in spatial features of ICNs, they do not fully evaluate the effect of age on other features, including those that may be frequency-specific.

While many rs-fMRI studies focus on low-frequency fluctuations of the BOLD signal up to 0.1 Hz (Biswal, Mennes et al. 2010), other studies using rs-fMRI and physiological recordings have established that multiple sub-bands of this spectrum up to 0.25 Hz provide meaningful information regarding neural processing (Wu, Gu et al. 2008, Song, Zhang et al. 2014). Most rs-fMRI studies typically use a repetition time (TR) of 2 seconds, resulting in the BOLD signal encompassing a range of frequencies between 0.01-0.25 Hz. Studies show that integration of ICNs differs across multiple bands of the frequency spectrum (Wu, Gu et al. 2008, Mather and Nga 2013, Gohel and Biswal 2015). Spectral power, amplitude of low frequency fluctuations (ALFF) and fractional amplitude of low frequency fluctuations (fALFF) are representative measures that take frequency into account. No previous studies have evaluated changes across multiple frequency bands throughout the lifespan in a comprehensive manner (Biswal, Mennes et al. 2010, Mather and Nga 2013, Gohel and Biswal 2015).

This study aims to replicate previous results of topological changes in ICNs across the lifespan using measures analogous to those previously used. We also aim to discern lifespan trajectories in multiple frequency bands. It is however important to note that this study is aimed at replicating the phenomenon of age-related changes in functional activation patterns and connectivity and not the methodology previously employed. We use IVA-L, an algorithmic extension of group ICA (GICA) that permits greater spatial variance in the estimated subject sources while making sure the sources are matched across subjects (Michael, Miller et al. 2013, Gopal, Miller et al. 2015, Laney, Westlake et al. 2015). We use previously identified measures ofspatial variability such as component volume (volume of independent components), location of the peak of functional clusters (Jolles, van Buchem et al. 2011, Gopal, Miller et al. 2016), and measures of temporal variability such as ALFF and fALFF over different bands to quantify variability across IVA-L derived independent sources (Zang, Yong et al. 2007, Zou, Zhu et al. 2008, Turner, Chen et al. 2012, Yan, Cheung et al. 2013) in addition to exploring age-related changes in functional connectivity. Some of these measures such as identifying the location of the peak of functional clusters, ALFF and fALFF in IVA-L separated sources have not previously been used to assess age-related changes over lifespan. Additionally, no known study has attempted to explore age-related changes in sub-bands of the frequency spectrum. With these additions and algorithmic changes, we aim to show that age-related changes in RSN ICNs are persistent conceptual phenomenon that can be elicited by using different approaches. We hypothesize that we will be able to reproduce previous results showing that ICN spatial patterns decrease in extent with age, and that functional connectivity in higher order networks exhibit regionally heterogeneous age-dependencies including linear and quadratic trajectories. We also expect to characterize novel age-related effects in the different frequency bands of the ICNs.

## Methods

### Participants

Onehundredandeighty-seven(agerange:6-85,56males/131females)right-handed participants’ data from the Nathan Kline Institute/Rockland Sample (http://fcon_1000.projets.nitric.org/indi/pro/nki.html, (Nooner, Colcombe et al. 2012)) were used after exclusion for diagnosed psychiatric disorders (no past or present DSM-IV diagnosis). Subjects with excessive head motion (> 3mm or 3 of motion in any direction) were excluded. The data were collected as per protocols approved by the institutional review board at NKI usinga 3T Seimens Trio scanner. Rs-fMRI data was collected for 10-minutes for each participant using a multi-band imaging sequence at TR = 1.4s, 2x2x2mm, 64 interleaved slices, TE = 30ms, flip angle = 65 , field of view = 224mm. A total of 404 volumes were collected.

### Data Processing

T1 images were brain extracted prior to post-hoc analysis using FSL’s BET algorithm. Resting state images were preprocessed using the DPABI toolbox (Yan, Wang et al. 2016) that employs FSL and SPM functions. Preprocessing included removing the first five volumes, realigning the images, co-registration to T1 structural images, smoothing using a 6mm Gaussian window from FSL, ICA-FIX to remove artifacts, and warping to the SPM EPI template (2mm resolution). The preprocessed rs-fMRI images were parsed using the IVA-L algorithm in the GIFT toolbox (http://mialab.mrn.org/software/gift/) into 75 independent components (IC) for each subject. A higher model order (IC = 75) allows for efficient estimation of individual RSNs without causing parts of different networks to be represented in the same component. This has been previously tested and compared to lower model orders in group ICA studies (Allen, Erhardt et al. 2012).

IVA-L is an extension of ICA that statistically identifies ICs in an input signal while maximizing mutual information across the subjects for a given IC (Kim, Attias et al. 2006, Kim, Attias et al. 2007, Lee, Lee et al. 2008). It uses a linear decomposition of sources similar to ICA but extends it to estimating the linear sources for each subject while maintaining the dependency across subjects to result in matched estimation as shown in equation 1.

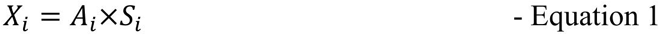

To improve computational efficiency of the algorithm, we used group principal component analysis (PCA) weights for each subject to initialize the IVA estimation. Group PCAweights have been typically used to initialize blind source separation of rs-fMRI data in group ICA implementations (Calhoun, Adali et al. 2001, Calhoun 2002, Calhoun 2004). Figure 1 shows a flowchart illustrating the data analytic pipeline used in the current study.

**Figure 1:**
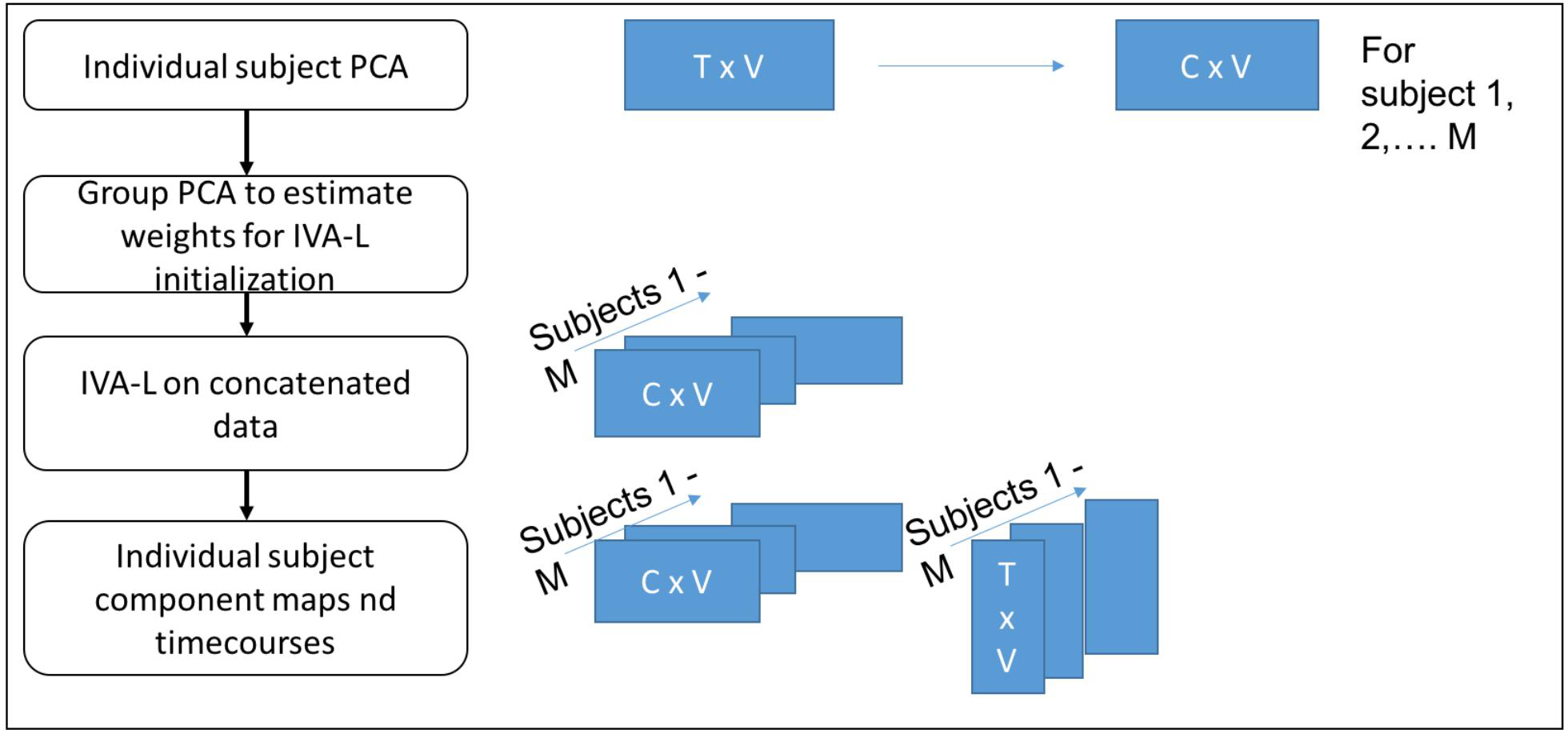
Flowchart representing the algorithmic implementation for data analysis using IVA-L

At the group level, ICs were visually inspected to identify non-artifactal (non-noise) components while those representing ringing, ventricles and other artifacts including speckling were removed. The individual subject component maps for these non-noise component were then z-scored. Non-noise IC timecourses were then despiked and detrended to remove drifts, and band-pass filtered (0.01 - 0.25Hz).

### Statistical Tests

#### a) Component Volume and Relationship with Age

For each subject’s non-noise IC, the component volume was computed as the number of voxels that survive a z-threshold of 3. We use this measure of the extent of the component’s functional cluster as an index of spatial variability in functional connectivity patterns across subjects. Component volume is dependent on the z-threshold chosen, but previous simulation studies have shown that reducing the z-threshold only increases the number of voxels that survive and does not affect the direction of the relationship (Gopal, Miller et al. 2016). Additionally, a z-threshold of 3 has been extensively used in previous studies and is sufficiently stringent to avoid false positives (Jolles, van Buchem et al. 2011, Woo, Krishnan et al. 2014). The component volume for each non-noise component was correlated with age and age^2^ across the participants and FDR corrected for multiple comparisons. A stepwise regression model was used to assess the amount of unique variance in the component volume explained by linear and quadratic interactions with age using SPSS v24 as shown in equations 2 and 3, respectively.

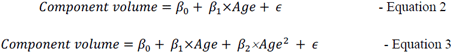

A significant change in F-statistic by adding the quadratic variable (age^2^) was considered a robust explanation of the quadratic relationship between age and component volume relationship. The sign of the  values represented the direction of correlation. The p-value for the linear regression model in Equation 2 had to be significant for the model to be identified as representative of the linear relationship between age and component volume. The p-values for both linear and quadratic models were FDR corrected.

#### b) Location of Cluster Peak and Relationship with Age

The location of the peak for each subject’s non-noise IC was identified and the distance of each subject’s peak from the centroid (estimated from all 187 subjects) was computed. The centroid was computed from the location of each subjects’ peak for a particular component over all subjects. This was used as an additional measure of spatial variance in rs-fMRI data. This distance was correlated with age across participants and FDR corrected for multiple comparisons.

#### c) Temporal Features

The post-processed non-noise IC timecourses from IVA-L were z-scored and the following tests were conducted. ALFF and fALFF were computed using the regressed timecourses for each subject and each component as per the following formulae.

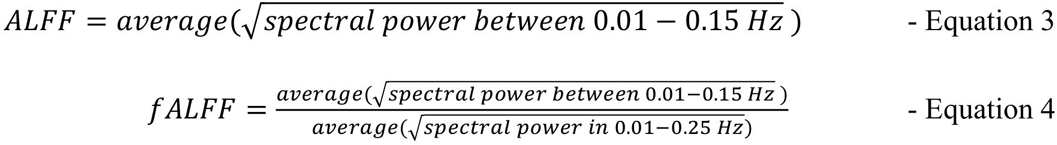

These overall ALFF and fALFF values were correlated with age for each non-noise IC component and corrected for multiple comparisons using FDR. Further inspection into bands ofthe frequencies was conducted to assess if different frequency bands exhibited different relationships with age. The spectra were sub-divided into four bins of the following frequency ranges - (0.01-0.027 Hz; 0.027-0.073 Hz; 0.073-0.198 Hz and 0.198-0.25Hz) as previously identified by multiple studies (Yue, Jia et al. 2015). The average square root power in each of these bins was computed and linear and quadratic relationships with age for each non-noise IC component were evaluated using a stepwise regression model as described in *a)* above. The p-values for these models were FDR corrected for multiple comparisons.

#### d) dStatic connectivity and relationship with age

For the non-noise IC components identified, static connectivity matrices (FNC matrices) were computed using correlations between IVA-L estimated timecourses. The non-noise IC components were ordered according to the ICNs they represented so as to clarify both within and between network connectivities in the matrix. The strength of connectivities between and within networks represented by the matrix were then modeled with age and age^2^ and FDR corrected for multiple comparisons to evaluate linear and quadratic changes in connectivity patterns with age using stepwise regression as previously described. The model fit (linear vs quadratic) was quantified using statistically significant change in F-statistic by adding the quadratic term to the linear regression model as described in *a)* above.

## Results

Visual inspection (by SGV and JN) of the 75 components yielded 29 non-noise IC components that were identified to represent known functional brain networks. These 29 components represent previously characterized functional networks in rs-fMRI data and are illustrated in Figure 2 (Biswal, Mennes et al. 2010).

### Component Volume and Age

Of the 29 components, the component volume of 20 components showed a negative linear relationship with age at an FDR threshold of p< 0.0140. These components were from the Frontal network, Salience network (SN), DMN, CEN, Auditory network, Visual network and sub-cortical structures/ Basal Ganglia. One component representing the SN showed a positive linear relationship with age. On further inspection, it was observed that this particular component represented parts of the thalamus in addition to the insular cortex. For these 21 components, the F-change after adding the quadratic age term to the linear model was not significant (p>0.05). Only one component representing the Basal Ganglia was found to have a negative quadratic relationship with age based on a significant F-change (*β*_*Lin*_= −0.193, *β_Quad_*= −0.329, *p_*Quad*_*= 0.000, *Change in F-stat*=22.504; *Change in R*^2^ = 0.133) strongest significant correlations (|r|>0.3) between component volume (at z-threshold of 3) and age as well as age^2^ are shown in Table 1, and Figure 3 shows the scatter plots for representative components’ volume from each network across age.

**Figure 2:**
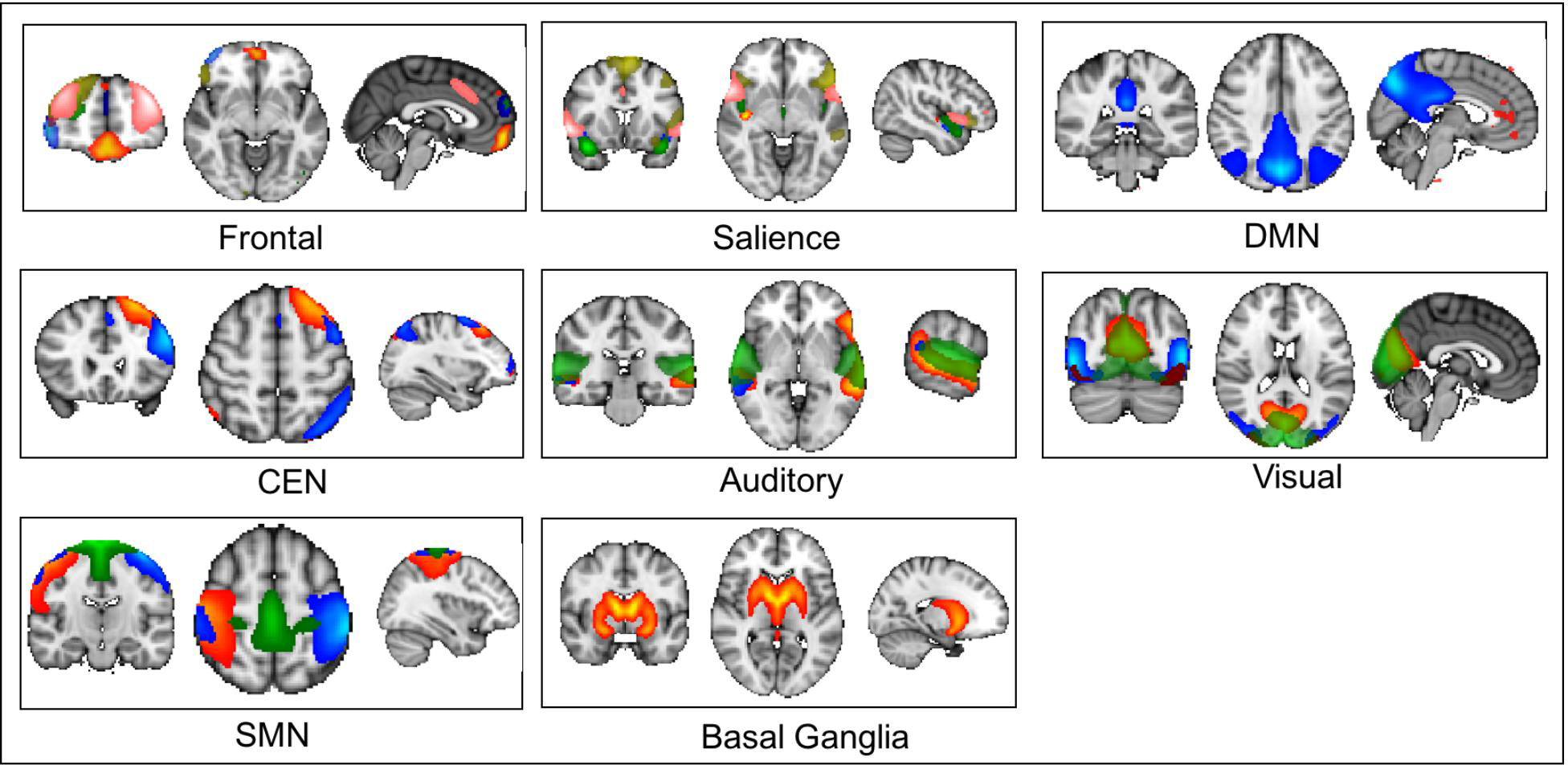
Spatial maps for all twenty-nine non-noise components that were used for further analysis.

**Figure 3:**
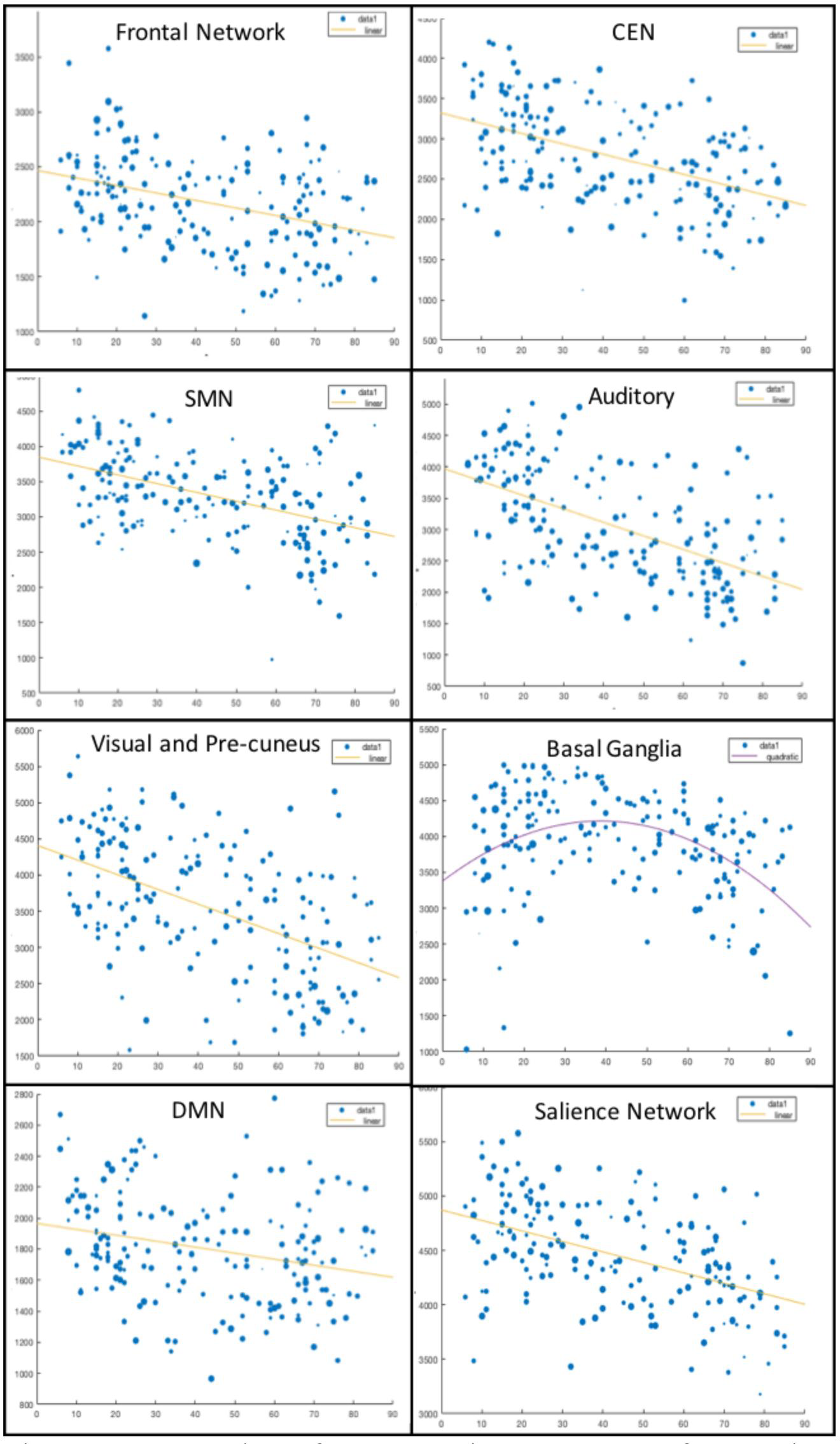
Scatter plots of representative components from each network with linear and quadratic relationship between component volume and age. The correlations and the corresponding p-values for each of the 22 components from each of the eight networks are shown in Table 1.

**Table 1:** Correlation of component volume at z-threshold = 3 and age and the corresponding p-value. The table presents components that had |r|>0.3 at an FDR corrected p<0.0140

| Comp Number | Label | Age |  |
| --- | --- | --- | --- |
|  |  | p_value | Correlation |
| 28 | FRONTAL | 0.00000 | -0.40856 |
| 30 | FRONTAL | 0.00001 | -0.31988 |
| 36 | FRONTAL | 0.00000 | -0.39472 |
| 39 | FRONTAL | 0.00000 | -0.37290 |
| 37 | Salience | 0.00468 | 0.20596 |
| 41 | Frontal | 0.00000 | -0.36782 |
| 58 | Salience | 0.00045 | -0.25424 |
| 64 | Salience | 0.00000 | -0.47920 |
| 21 | DMN | 0.00022 | -0.26712 |
| 33 | Frontal | 0.00003 | -0.30231 |
| 49 | Auditory | 0.00000 | -0.47915 |
| 56 | CEN | 0.01264 | -0.18207 |
| 46 | Auditory | 0.00000 | -0.55784 |
| 54 | Auditory | 0.00000 | -0.52219 |
| 50 | Visual | 0.00000 | -0.51309 |
| 62 | Visual | 0.00000 | -0.48600 |
| 69 | Visual | 0.01178 | -0.18384 |
| 55 | SMN | 0.00000 | -0.46700 |
| 59 | SMN | 0.00000 | -0.39120 |
| 63 | SMN | 0.00000 | -0.35755 |

### Location of Cluster Peak and its Relationship with Age

The distance of component cluster peaks from the group centroid were not significantly related to age for any of the 29 non-noise IC components. This suggests that there was no significant age-related variance in functional localization of clusters for this dataset.

### Temporal Features

Of the 29 non-noise IC components, 2 components showed statistically significant negative correlations between overall ALFF and age at an FDR corrected threshold of p< 0.0038. Table 2 shows the correlation r and p values for these components. For overall fALFF, 16 componentsshowed statistically significant negative correlations with age at FDR corrected p<0.0239 whose r and p values are also shown in Table 2.

**Table 2:** Components with statistically significant correlation between fALFF/ ALFF and age at FDR corrected p-value< 0.0239 (fALFF) and 0.00198 (ALFF).

|  | fALFF |  | ALFF |  |
| --- | --- | --- | --- | --- |
|  | p value | correlation | p value | correlation |
| 1 | 0.0000010 | -0.34940 |  |  |
| 2 | 0.0052792 | -0.20322 |  |  |
| 3 | 0.0000009 | -0.35086 |  |  |
| 4 | 0.0001083 | -0.27932 |  |  |
| 5 | 0.0000001 | -0.37931 |  |  |
| 7 | 0.0000011 | -0.34725 |  |  |
| 8 | 0.0000000 | -0.40007 |  |  |
| 9 | 0.0000000 | -0.41868 |  |  |
| 10 | 0.0000368 | -0.29695 |  |  |
| 13 | 0.0000003 | -0.36445 |  |  |
| 14 | 0.0000106 | -0.31589 |  |  |
| 16 | 0.0001343 | -0.27565 |  |  |
| 17 | 0.0238517 | -0.16520 |  |  |
| 19 | 0.0003561 | -0.25840 |  |  |
| 22 | 0.0065753 | -0.19809 |  |  |
| 24 |  |  | 0.00075 | -0.2444 |
| 26 | 0.0128955 | -0.18155 |  |  |
| 27 |  |  | 0.002 | -0.2441 |

The binned spectral power values (binned mean of squared root of spectral power of frequencies included in each bin) showed novel results in terms of relationship between activation in different frequency bands and age. We found that the spectral power in bins 1 and 2 linearly decreased with age while the spectral power in bins 3 and 4 linearly increased with age. Additionally, band specific relationship with age were found in specific ICNs. Some of the interesting observations included – Bin 1 (0-0.027 Hz) - spectral power in only the SN components showed a linear decline with age (FDR corrected), Bin 4 (0.198 – 0.25) spectral power in only the frontal network showed a linear increase with age, the DMN components only showed linear decline in Bin 2 (0.027-0.073) with age, the SMN showed no linear or quadratic relationships with age and the most number of components with age relationships were in Bins 2 (0.027-0.073 Hz) and 3 (0.073 – 0.198 Hz).

### Static connectivity and relationship with age

In line with previous results, we found a distribution of positive and negative linear and quadratic relationships between connectivity strengths and age of different ICNs. While most correlations did not survive FDR correction (p<0.0011), we found that SN connectivity with the DMN had a positive linear relationship with age, but SN connectivity with the sensorimotor network showed a negative linear relationship age. Additionally, CEN connectivity with the sensorimotor network showed a negative linear relationship with age along with connectivity between sensorimotor network components (within network). In addition, quadratic relationships were found between SN and visual network components (positive) and sensorimotor andauditory components (negative). These connectivities and their relationship with age are represented in Figure 5.

**Figure 4:**
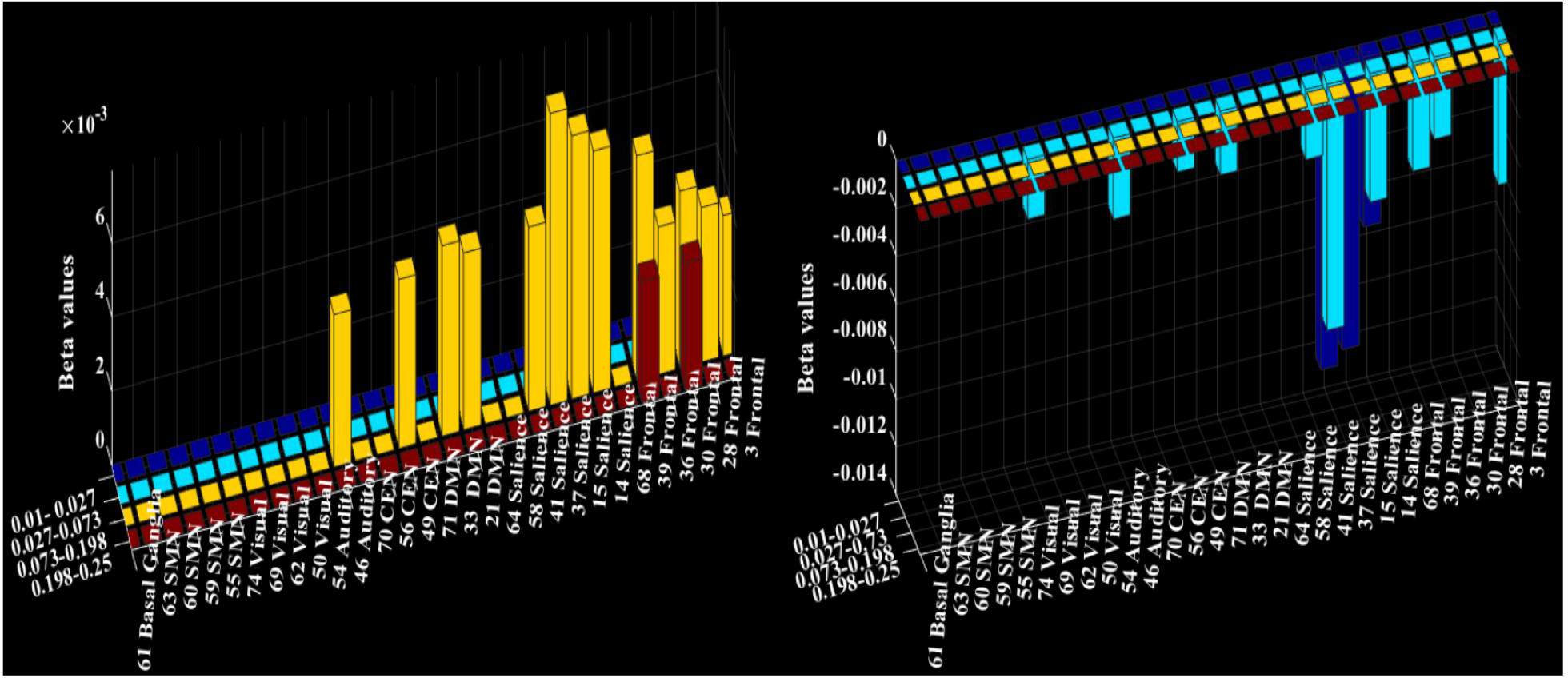
Beta values of binned spectral power modelled with age that survived FDR correction at p<0.0088. The left panel shows the components with positive linear relationship with age and the right panel shows the components with negative linear relationship with age. It can be clearly see from these figures above that a linear increase in spectral power was observed with age in the higher frequency bands i.e. Bin 3 and Bin 4 whereas a linear decrease in spectral power was observed with age in the lower frequency bands i.e. Bin 1 and Bin 2.

**Figure 5:**
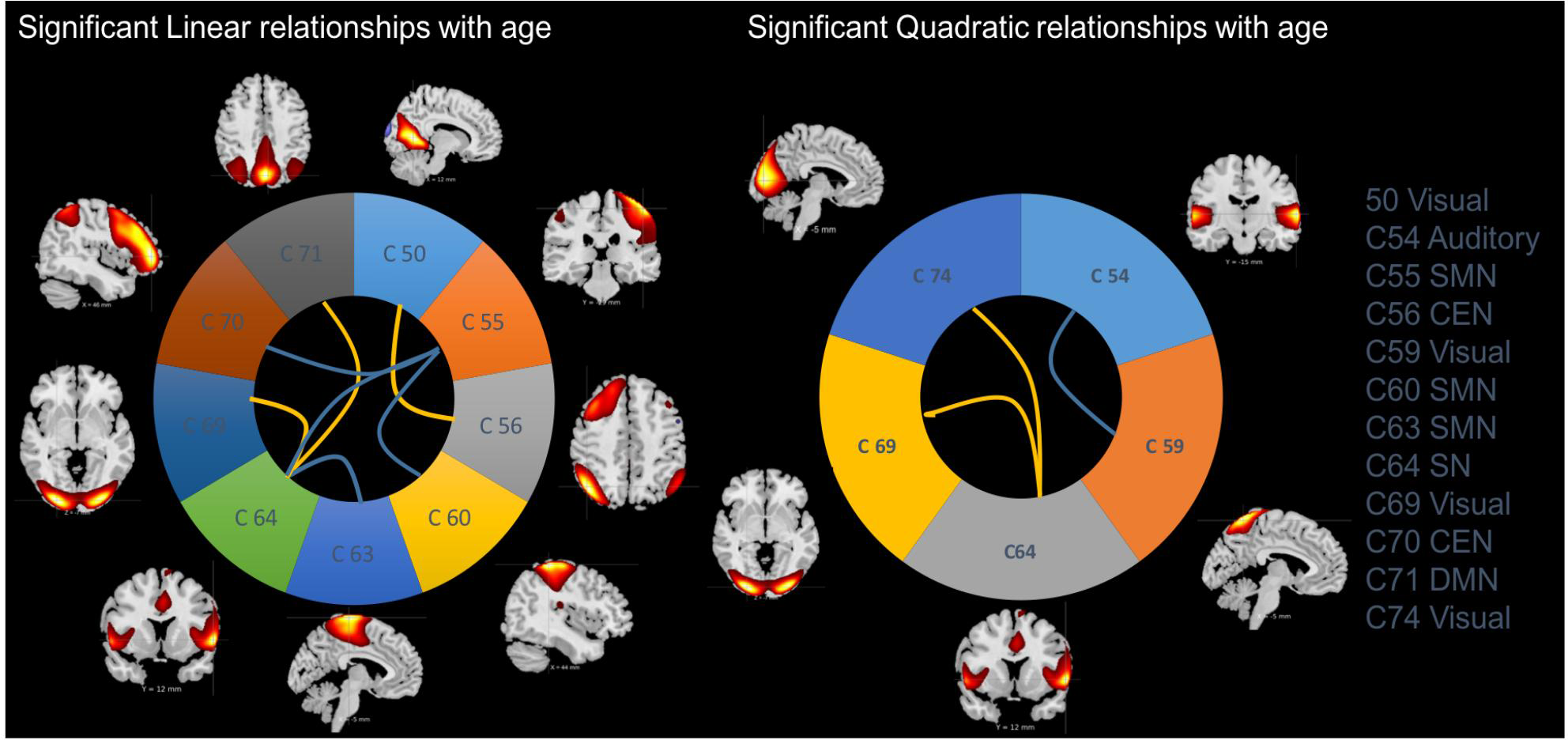
Static FNC correlation with age. The left panel shows the linear relationship between network connectivities and age while the right panel shows the quadratic relationship with age. The lines represent the pair of components whose correlation with age/age2 is significant. The blue lines represent positive relationship with age and the yellow lines represent negative relationship with age. For e.g. in the left panel, connectivity between C54 and C64 has a positive linear relationship with age and in the right panel, connectivity between C59 and C54 has a positive quadratic relationship with age. These relationships are FDR corrected with p<0.0011.

## Discussion

This study validates previously identified age-related topological changes in the human functional connectome across the lifespan (6-85 years) as well as provides novel insights into the behavior of different networks in different frequency bands (Betzel, Byrge et al. 2014, Cao, Wang et al. 2014, Yang, Chang et al. 2014, Gohel and Biswal 2015). We employed IVA-L, an innovative whole brain multivariate analysis technique to extract ICNs from rs-fMRI data, and validation and exploration of age-related changes. We found functional changes across the lifespan that direct our attention to the increasing stability of brain networks with age. We show that age-related variance primarily exists in the spatial extent of the cluster of ICNs with the functional localization remaining constant. Additionally, age-related variance in functional connectivity was found in ICNs involved in higher order cognitive processing. Additionally, we found that activation in slower frequency bands decreased with age while faster frequency band activation increased with age. These findings demonstrate that age-related changes in ICNs extend beyond topographic features and connectivity.

### IVA-L vs GICA

Studies of healthy children and the elderly have consistently used GICA as a method to assess age-related changes in functional connectivity (Huang, Hsieh et al. 2015, Muetzel, Blanken et al. 2016, Sole-Padulles, Castro-Fornieles et al. 2016). However, there are some limitations in using this algorithm to explore spatial variability in ICN features. Algorithmically, GICA might lead to loss of some variability in spatial features due to the two PCA steps for dimensionality reduction. Additionally, GICA estimates a set of group components shared by allsubjects from which each subjects’ individual component maps and timecourses are back-reconstructed. This further constrains the individual variability in spatial and temporal features that can be incorporated. Studies have shown that algorithmic modifications and extensions of ICA exist that improve the inter-subject variability incorporated in individual component maps and timecourses (Du, Li et al. 2011, Cao, Wang et al. 2014, Michael, Miller et al. 2014, Silva, Plis et al. 2014). IVA is one such algorithm that overcomes the limitations of GICA by applying a single PCA to the data for reduction of dimensionality and also estimating the sources for each subject separately. IVA estimates sources that are independent from each other but establishes a dependency between the subjects that allows for the source components to be matched across subjects. This dependency however, is different from forcing all the subjects to share source components, thereby allowing for greater variability to be incorporated. While other algorithms and modifications of ICA exist that attempt to address these concerns, we choose to use the IVA-L algorithm based on previous evidence from simulation studies and applications to schizophrenia that demonstrate its effectiveness in assimilating inter-subject variability (Kim, Attias et al. 2006, Lee, Lee et al. 2008, Michael, Miller et al. 2014, Gopal, Miller et al. 2015, Silva, Plis et al. 2014).

### Topographic changes in ICNs

Functional networks are affected by factors such as age, sex and neuropsychological disorders (Biswal, Mennes et al. 2010). In infancy and early age i.e. 6 – 10 years, Gao et. al., Muetzel et. al. and de Bie et. al. show that the default mode network is present but immature (de Bie, Boersma M Fau - Adriaanse et al. 2012, Gao, Alcauter et al. 2015, Muetzel, Blanken et al. 2016). On the other hand, Huang et. al. show that age-related decline in functional connectivity exists in older participants 51 – 85 years of age (Huang, Hsieh et al. 2015). These studies showthat a large-scale network structure is present from infancy, and that this evolves with age [Jolles, Muetzel,]. Jolles et. al used group ICA to explore age-related variability in topological features of blind source separated components. They found that ICNs show a decrease in the spatial extent of component clusters from childhood to young adulthood indicating a more “widespread” network architecture in childhood.

Our analysis also found that component volume (i.e. extent of IVA-L separated clusters) was negatively correlated with age. Reduction in component volume is directly related to reduced cortical activation with age, indicating an increased stability in functional connectivity patterns across the lifespan. This corroborates results from task-based studies that posit that functional activation patterns transition from diffuse to focal with age (Durston, Davidson et al. 2006). In conjunction with the finding of no significant variation in the location of the clusters peak location, we can surmise that the spatial variability observed is primarily in the extent of the clusters and that functional localization does not change with age. Furthermore, studies suggest that ICNs transform from focal to distributed over the lifespan, indicating that as networks become more specialized, they tend to strengthen long-range communications with other specialized networks (Betzel, Byrge et al. 2014, Cao, Wang et al. 2014). The previous work by Betzel and Cao used modularity and local/global efficiency to assess lifespan changes and showed that changes in functional connections in the brain were distance-dependent. Our study generally corroborates findings that ICNs transition from diffuse to focal over the lifespan.

It is noteworthy that this age-related reduction in component volume was consistently linear across most networks. This indicates that a general and widespread decrease in cortical employment is associated with maturation of ICNs. However, the Basal Ganglia network component that included the thalamus showed a quadratic decline with age, suggesting a morenuanced developmental trajectory for these subcortical regions. The one SN component with representation in thalamus showed a positive linear relationship with age, which might be attributed to the functional activation in the thalamus. These findings are broadly consistent with previous reports of increased ICN segregation with age.

### Functional connectivity changes between ICNs

Functional connectivity within and between ICNs has been extensively studied using multiple methods including ICA, graph-theoretical approaches as well as whole-brain clustering approaches(Betzel, Byrge et al. 2014, Cao, Wang et al. 2014, Yang, Chang et al. 2014). These studies consistently reveal that functional connections within ICNs weaken with age while connections between ICNs increases. Additionally, these changes can be selective to ICNs involved in higher order cognitive processing such as DMN, Salience, and central executive network (Uddin 2011). Betzel et. al show patterns of increasing connectivity between Salience network and DMN and other higher order cognitive networks (Betzel, Byrge et al. 2014) with age. Yang et. al were further able to differentiate age-related changes in sub-parts of the DMN (Yang, Chang et al. 2014). Jolles et. al. showed that majority of networks showed regional decreases in functional connectivity such that within network connectivity was decreased in children compared with adults (Jolles, van Buchem et al. 2011). Our results do not contradict the general trend of age-related changes in functional connectivity found by these studies. We show that functional connectivity between the SN and the DMN, SN and the visual network and CEN and visual network (precuneus) linearly strengthens with age. On the other hand, we also found that connectivity between the SN and SMN and CEN and SMN linearly decreased with age. We also found that within network connectivity in the SMN decreased linearly with age. Moreover, quadratic increase in the connectivity between salience and visual network and decline in connectivity between SMN and auditory network with age were found.

While we verify that functional connectivity between higher order networks strengthen with age, there are some interesting aspects to note. The SN is known as a hub that mediates information flow between other networks involved in higher order cognition and information processing (Uddin, Supekar et al. 2010, Uddin 2015). It has also been shown that structural integrity of the salience network is required to modulate the function of the DMN and attention networks (Buckner, Andrews-Hanna et al. 2008). We found significant age-related linear and quadratic changes in the connectivity of the SN with the DMN, SMN and visual network components including precuneus, implying that age-related changes in the functioning of DMN, SMN and visual network can be driven by changes in the functional integrity of the SN itself.

### Spectral relationship changes in ICNs

Lifespan studies have evaluated age-related changes in low frequency fluctuations in BOLD fMRI signals using ALFF and fALFF (Biswal, Mennes et al. 2010, Hu, Chao et al. 2014). These measure characterize the spontaneous fluctuation in the BOLD signal and provide us with a meaningful information regarding dynamic neural activations. Biswal et. al. found that medial wall structures showed significant age-related decreases in fALFF. Hu et al. (Hu, Chao et al. 2014) showed consistent results, suggesting that age related decline in fALFF could suggest vulnerability of networks with aging. However, these previous studies used measures of low-frequency fluctuations in fMRI data prior to applying whole-brain source separation algorithms. While such an approach is advantageous in identifying systemic changes in ALFF and fALFF, using these measures in ICA/IVA separated source timecourses allows us to assess age-related changes at the level of ICN integration while also eliminating potential sources of noise.

The current study used ALFF and fALFF to characterize changes in amplitude of low frequency fluctuations of ICN timecourses estimated by IVA-L. Our results show an age-related decline in both ALFF and fALFF. While ALFF is a more stringent measure of spontaneous neural activations, we only found SMN and visual network decline with age. In contrast fALFF illustrated a widespread decline with age over the frontal network, salience network, DMN, auditory network and the visual network. These results verify and extend previous results in novel ways including the application of IVA-L. The range of BOLD signals used to assess low frequency fluctuations using ALFF and fALFF typically include 0.01-0.1 Hz for ALFF and 0.01-0.25 Hz for fALFF. This range of frequencies includes multiple bands that might relate to demographic and behavioral features differently.

Few studies have explored age-related changes in ALFF across frequency bands (Mennes, Kelly et al. 2010, Mather and Nga 2013). Mather and Nga showed that different frequency bands had different relationships with age in the thalamus. They showed that averaged fALFF within the thalamus shifts in directionality; the band 0.01-0.10 Hz showed negative correlations with age, 0.10-0.27 Hz showed positive correlations with age, and 0.198-0.25 Hz showed negative correlations with age. Most relevant to our hypothesis, Gohel et. al. evaluated the functional integration of networks in different frequency bands (slow – 5: 0.01-0.027 Hz, slow -4: 0.027-0.073 Hz, slow -3: 0.073 – 0.198 Hz, slow -2: 0.198-0.5 Hz and slow -1: 0.5-0.75 Hz)(Gohel and Biswal 2015). They used GICA to estimate the ICNs in the different frequency bands of the fMRI data and found that the large-scale network architecture was consistent across all frequency bands but the slow -3 and slow -4 bands encompassed most of the power compared to other frequency bands across ICNs. The hypothesis that different frequency bands relate differently to age is further bolstered by electrophysiological studies. However, no known studyhas evaluated age-related changes in low frequency fluctuations in a large range of frequency bands. Results of the current study further confirm that network modulation with age is frequency dependent.

Our results show that spectral power in the 0.01-0.073 Hz band demonstrates a negative relationship with age, whereas the spectral power in 0.073-0.25 Hz band showed a positive relationship with age. The slowest (0.01-0.027 Hz) and fastest (0.198- 0.25 Hz) frequency bands showed significant linear trends with age in the SN and the frontal network components, respectively. Moreover, most of the age-related changes in the four bands we studied were observed in the SN and frontal network components. Also, DMN showed age-related changes only in the slow -3 bin (0.073-0.198 Hz) reflecting that higher frequency activation in the DMN increased with age. These findings are in-line with the hypothesis that with age, higher order cognitive processes become more developed. It is noteworthy that the SMN components did not show age-related changes in the spectral power of any of these frequency bands. These results not only highlight the importance of exploring signals in multiple frequency bands but also direct our attention to the fact that age-related increases in higher frequency activation across components may support cognitive maturation. The observed decrease in lower frequency activation could relate to networks becoming increasingly differentiated and stable with age.

## Conclusion

The characterization of lifespan evolution of ICNs has been of immense interest to researchers focusing on typical and atypical developmental trajectories. Here we investigate cortical employment and functional localization, reproducing findings that ICNs employ less cortical surface as they specialize with age. We show that the salience network plays a critical role in age-related changes, as represented by changes in coupling with other networks. Lastly, we evaluate spontaneous neural activations in multiple frequency bands and show that age-related changes in ICNs differ across different frequencies. Quantifying the variability in spatial and temporal features of ICNs across the lifespan will allow us to not only characterize typical developmental trajectories, but also help in identifying abnormalities in these trajectories associated with neuropsychological disorders.

